# Longitudinal hair cortisol in bipolar disorder and a mechanism based on HPA dynamics

**DOI:** 10.1101/2023.07.03.546860

**Authors:** Tomer Milo, Lior Maimon, Ben Cohen, Dafna Haran, Dror Segman, Tamar Danon, Anat Bren, Avi Mayo, Gadi Cohen Rappaport, Melvin McInnis, Uri Alon

## Abstract

Bipolar disorder (BD) is a dynamic disease in which mania, depression and mixed states vary on a timescale of months to years. BD patients characteristically exhibit elevated levels of the hormone cortisol. Persistently elevated cortisol can also cause mood episodes in a substantial fraction of the general population. Although BD is a dynamic disease that is related to cortisol, longitudinal cortisol dynamics in BD have rarely been studied. Here we use hair to measure past cortisol where each cm of hair corresponds to a month of growth. Cortisol was measured in 12 cm hair samples from people with BD (n=26) and controls (n=59), corresponding to one year of cortisol data. We found that hair cortisol exhibited a frequency spectrum with enhanced year-scale fluctuations whose amplitude was about 4-fold higher on average in BD compared to controls. Cortisol in the proximal 2 cm hair segment correlated with mood scales that report on mood in the past two months. In line with the notion that cortisol correlates with mood, we find that the mean frequency spectrum of depression (n=266) and mania (n=273) scores from a large longitudinal study of BD is similar to the hair cortisol spectrum from the present cohort. Taken together, these results suggest a mechanism for BD as the intersection of two neuropsychological traits: cortisol-induced mood episodes (CIM) and high emotional reactivity (ER). High ER causes fluctuations in which cortisol is elevated for months, as shown by a mathematical model of the hypothalamic-pituitary-adrenal (HPA) axis that regulates cortisol. In individuals with CIM, the magnitude of these persistent cortisol fluctuations can be high enough to trigger mood episodes. Thus, this study combines longitudinal cortisol measurements and mathematical modeling to provide a potential mechanistic link between the timescales of cortisol and moods in BD.

## Introduction

Bipolar disorder (BD) is a mood disorder with temporal variation between mania, depression, mixed states and euthymic states. The biological mechanisms underlying BD are heterogeneous and complex (Vieta et al. 2018; Carvalho, Firth, and Vieta 2020). Advances in understanding the pathophysiological mechanisms of BD are needed in order to develop new avenues for treatment (Bauer and Dinan 2015; Trinetti, Sforzini, and Pariante 2021).

A defining characteristic of the mood changes in BD is their timescale of months to years. Mania episodes typically last several weeks. Depression episodes typically last months or longer (Solomon et al. 2010). There does not appear to be a typical frequency for mood changes (Cochran et al. 2018), but rather a wide range of frequencies with fluctuations on the scale of months to years. Episode rate can range from four times or more per year in rapid-cycling BD to once every few years (Lee et al. 2010; Nierenberg et al. 2010; Wells et al. 2010), with some people experiencing euthymic conditions without aberrant mood swings most of the time. The origin(s) of these long timescales and the underlying factors contributing to their temporal variability is not understood.

The purpose of this study was to identify potential mechanisms contributing to the observed timescales of bipolar disorder. To do so, we focused on a physiological system implicated in bipolar disorder -the HPA axis which controls the stress hormone cortisol (Daban et al. 2005; Watson and Mackin 2006). Cortisol is produced by the adrenal cortex, under control of ACTH from the pituitary, which in turn is induced by CRH secreted from the hypothalamus in response to stress inputs. Cortisol has wide-ranging effects on metabolism and cognition (Kim and Kim 2023; Brillon et al. 1995; de Souza-Talarico et al. 2011). It has receptors in many brain areas including the prefrontal cortex and hippocampus (Dedovic et al. 2009). The effects of cortisol depend on whether the stress is acute or chronic. Chronically high levels of cortisol are generally damaging to neurological and physiological systems (Lupien et al. 2007; Cox et al. 2015).

Recent advances in mathematical modeling show that the HPA glands, the pituitary corticotrophs and the adrenal cortex, can grow or shrink on the timescale of months, providing fluctuations in cortisol that can last months to years. These gland-mass changes arise from the fact that HPA hormones CRH and ACTH are growth factors for the secretory cells in the pituitary and the adrenal cortex, respectively (Karin et al. 2020; Karin, Raz, and Alon 2021; Tendler et al. 2021). The timescale for mass changes of these glands is determined by the turnover time of their cells, which occurs on the order of 1 to 2 months (Karin et al. 2020). These growth-factor interactions cause the pituitary and adrenal masses to act as a damped oscillator, which can be induced by stress inputs to exhibit noisy oscillations of gland masses with a wide range of frequencies corresponding to periods of months to years (Maimon et al. 2020). Indeed, a control population without BD showed months-scale fluctuations of hair cortisol, with the strongest contributions from the lowest frequency measured, corresponding to a period of one year (Maimon et al. 2020).

Further implicating the HPA axis in BD are the collective studies showing the average serum, saliva and hair cortisol levels are about two-fold higher in BD patients than in control populations (van den Berg et al. 2020; Coello et al. 2019; Herane-Vives et al. 2020; Koumantarou Malisiova et al. 2021; Streit et al. 2016; Belvederi Murri et al. 2016). BD patients also exhibit dysregulation of the HPA axis, measured as a reduced suppression of cortisol in the dexamethasone test (Rush et al. 1997; Rybakowski and Twardowska 1999; Schmider et al. 1995; Watson et al. 2004) and altered pituitary size (Delvecchio et al. 2018; MacMaster et al. 2008; Takahashi et al. 2010).

Elevated cortisol is not only a feature of BD, but can also cause BD-like mood episodes. People treated with glucocorticoid steroids, which are cortisol analogues, for a variety of ailments are at high risk to develop manic or depressive symptoms (L. Fardet et al. 2007; Laurence Fardet, Petersen, and Nazareth 2012; Judd et al. 2014). The risk for developing these neuropsychiatric symptoms increases with glucocorticoid dose (Laurence Fardet, Petersen, and Nazareth 2012), and a previous history of unstable mood patterns further increases the risk (Judd et al. 2014).

Similar findings occur in Cushing’s syndrome, in which cortisol levels are elevated due to hormone-secreting tumors (Pivonello et al. 2015; Sonino and Fava 2001). About 30% of Cushing patients experience mania or hypomania, and 50-60% present with depression (Sonino et al. 1998). One study found that of 30 patients with Cushing’s syndrome, 20 met criteria for depressive episode and 8 of these also reported an episode of mania or hypomania during their endocrine disturbance (Haskett 1985). These findings suggest that a fraction of individuals are sensitive to chronically high cortisol levels in triggering mood episodes.

Several additional lines of evidence suggest that the HPA axis plays a role in the pathophysiology of BD. Early BD episodes are often triggered by stressful life events (Alloy et al. 2005, 200; Bender and Alloy 2011). Unaffected relatives of patients with BD display abnormal HPA axis activity (Ellenbogen et al. 2006; Krieg et al. 2001), though this is not an endophenotype of BD, but rather seems related to environmental risk factors, such as childhood trauma (Belvederi Murri et al. 2016).

Despite the fact that BD is a dynamic disease with strong ties to cortisol, the dynamics of cortisol in BD over the timescales of months to a year is poorly characterized. Previous studies of BD typically measured cortisol at one or two time points (Streit et al. 2016; van den Berg et al. 2020; Koumantarou Malisiova et al. 2021). There is thus a need for longitudinal cortisol measurements over months in a cohort of people with BD.

Here we addressed this by using hair to measure longitudinal cortisol levels in a BD population and a control population over one year. We compared the cortisol dynamics to the dynamics of mood scales (McInnis et al. 2018). The results were analyzed by using the gland-mass mathematical model of the HPA axis. Informed by these findings, we propose a mechanism for the timescales of BD based on the HPA axis.

## Results

### Longitudinal hair cortisol in participants with BD is higher than in controls

We compared hair cortisol from participants with BD (n=26) to control participants (n=59). We measured hair cortisol in 12 cm of hair, corresponding to about a year of growth. Cortisol was evaluated in 6 segments of 2 cm each. Cortisol levels declined along the length of hair as previously described (Maimon et al. 2020). In all six hair segments, measurements in participants with BD had about 2-fold higher median cortisol than control participants (Figure 1).

**Figure 1.**
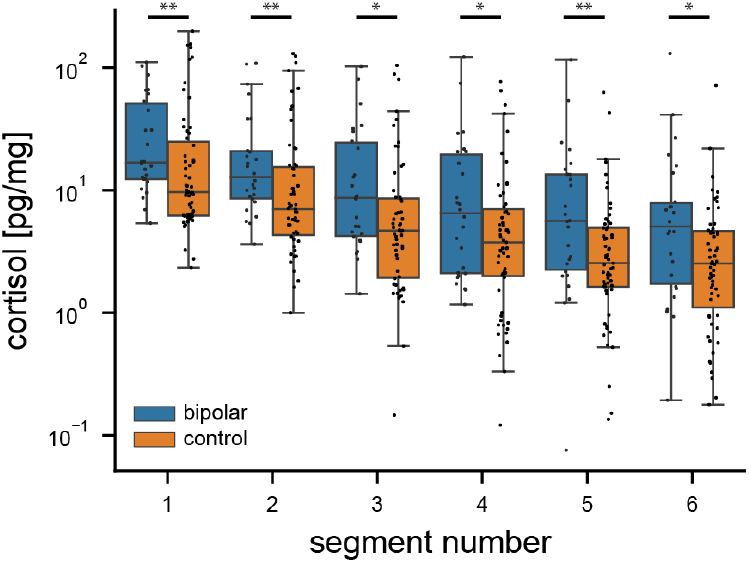
BD participants had higher hair cortisol than control participants. Cortisol levels of control (orange) and BD patients (blue) along six successive 2-cm hair segments of a 12cm hair sample, where segment 1 is closest to the scalp. Effect sizes, calculated as the ratio between BD and control median cortisol, and Mann-Whitney U test p-values for segment 1 to 6 are: 1.7, p=0.008; 1.8, p=0.003; 1.85, p=0.01; 1.7, p=0.04; 2.2, p=0.005; 2, p=0.01.

### BD participants had enhanced year-scale fluctuations in cortisol levels

We next studied the variation of cortisol over time in each individual. We accounted for the natural decline along the hair and evaluated the cortisol fluctuations in both control and BD participants using the methods developed in (Maimon et al. 2020). This resulted in normalized cortisol readings for six 2cm segments (representing 2 months of growth) points from each 12 cm hair sample (Figure 2A).

**Figure 2.**
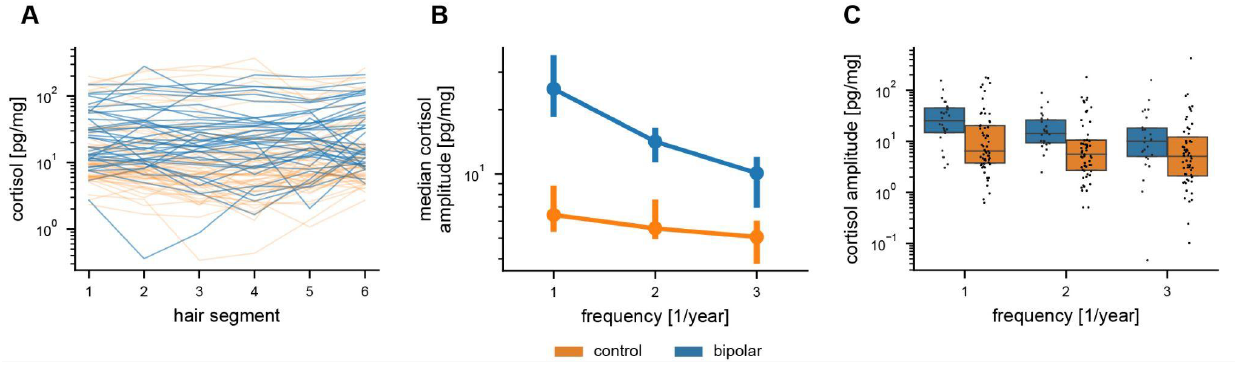
Participants with BD showed enhanced year-scale fluctuations of hair cortisol. **(A)** Individual cortisol time series of 6 hair segments after correcting for the cortisol decay along the hair. **(B)** Median Fourier amplitudes of longitudinal hair cortisol. Error bars are 68% confidence intervals from bootstrapping. The ratios between BD and control median fourier amplitudes were: 3.9 (p=0.001) for 1 *year*^−1^ frequency; 2.55 (p=0.0003) for 2 *year*^−1^ frequency, and 2 (p=0.02) for 3 *year*^−1^ frequency. The ratio between the median amplitude of the lowest and the highest frequencies is 2.5 (p=0.006) in BD and 1.3 (p=0.02) in controls. All p-values from Mann-Whitney U test. **(C)** Fourier amplitudes of all participants (dots). Boxes show 25%, 50% (median) and 75% percentiles.

To evaluate the contribution of different timescales to the cortisol dynamics we used Fourier analysis. Fourier analysis decomposes the signal into a sum of oscillatory components at different frequencies, and quantifies the contribution of each frequency component as a Fourier amplitude. Six segments, spanning one year, allow three frequencies to be detected: 1 *year*^−1^, 2 *year*^−1^, and 3 *year*^−1^, representing periods of a year, 6 months, and 4 months, respectively.

In both groups the Fourier amplitudes declined with frequency (Figure 2B-C), in agreement with (Maimon et al. 2020). In controls, median amplitude at the lowest frequency (1 *year*^−1^) was about 1.3 (p=0.02, Mann-Whitney U test) times higher than the median amplitude at the highest frequency (3 *year*^−1^). In the BD group, the median amplitude at the lowest frequency was higher by a factor of about 2.5 (p=0.006, Mann-Whitney U test) than that of the highest frequency.

The Fourier amplitudes of the BD group were significantly higher than controls (p=0.001, 0.0003, 0.02 for the three frequencies, Mann-Whitney U test). At the lowest frequency, the median amplitude is 4 times larger in BD than in controls. This shows that BD participants exhibit year-scale peaks and troughs in cortisol levels which are much larger than those of the control group.

### Hair cortisol of participants with BD correlated with depression and anxiety mood scores

We also tested whether hair cortisol was associated with mood scales. At the time the BD participants contributed the 12cm hair sample, we evaluated them with four mood scales, the YMRS for mania, BDI and HDRS for depression and HAM-A for anxiety (see Methods). Thirteen participants returned and donated a second hair sample more than two months after the first sample, and were evaluated using these mood scales a second time. One participant did not fill out the scales. In total we had 38 hair samples with 38 corresponding sets of mood scores.

Since the mood scales are designed to estimate mood over the recent weeks, we compared the mood scale scores to the 2 cm segment of hair most proximal to the scalp, which corresponds to cortisol in the 2 months prior to hair collection.

Low YMRS mania scale scores were recorded for all participants (mean score=2, std=3.3), indicating that none of the participants showed significant manic symptoms during the study. YMRS scores did not correlate with log cortisol (*r* = 0. 23, *p* = 0. 26, adjusted partial Pearson correlation).

A wide range of scores were recorded in the depression and anxiety scales, indicating a range of depression and anxiety symptoms in the BD group. The two depression scales BDI and HDRS both correlated positively with log hair cortisol (*r* = 0. 47, *p* = 0. 007 BDI, *r* = 0. 55, *p* = 0. 001 HDRS, adjusted partial Pearson correlation). The anxiety scale HAM-A also correlated positively with log cortisol (*r* = 0. 45, *p* = 0. 01, adjusted partial Pearson correlation). We also repeated the analysis excluding the 13 repeat measurements. The HDRS correlation remained significant (*r* = 0. 53, *p* = 0. 025). The other two scales show insignificant positive correlation trends (BDI *r* = 0. 38, *p* = 0. 09 and HAM-A *r* = 0. 3, *p* = 0. 15).

The BDI, HAM-A and HDRS scale scores also correlated well with each other (*r* = 0. 88, *p* = 6⋅10^−13^ BDI vs. HDRS, *r* = 0. 72, *p* = 5⋅10^−7^ BDI vs. HAM-A, *r* = 0. 66, *p* = 7⋅10^−6^ HDRS vs. HAM-A) indicating satisfactory reliability of the questionnaires. We conclude that hair cortisol in the 2cms most proximal to the scalp correlates with depression and anxiety mood scales.

### A longitudinal BD mood scale study show similar frequency spectra to hair cortisol

We next assessed whether the low frequency components observed in the hair cortisol dynamics of BD participants resemble the frequencies found in a much larger study of longitudinal mood measurements. For this purpose we analyzed longitudinal mood scale scores data from the Prechter BD cohort, where participants complete the PHQ-9 and the ASRM questionnaires every two months over multiple years (McInnis et al. 2018) (see Methods).

The Prechter BD cohort includes 541 bipolar type-1 patients and 267 controls. We analyzed the BD participants with at least two years of consecutive mood measurements and took their longest consecutive time series for each mood scale. Thus, we included 266 BD and 179 control time series of depression scores (PHQ-9), and 273 BD and 178 control time series of mania scores (ASRM). In total we analyzed 24,627 mood measurements.

To estimate the Fourier frequency spectrum of the mood time series we used the Lomb-Scargle method (VanderPlas 2018; Lomb 1976; Scargle 1982) which is suitable for time series of differing lengths. The PHQ-9 exhibited a Fourier spectrum displaying the highest amplitudes at low frequencies (Figure 3A). The 1 *year*^−1^ median amplitude was 1.2-fold (p=0.01, Mann-Whitney U test) higher than the 3 year^−1^ median amplitude. The Fourier amplitudes were higher in BD participants than in controls. A similar feature was observed in the ASRM scores with declining frequency amplitudes along the Fourier spectrum, where the 1 *year*^−1^ median amplitude was 1.4-fold (p=0.001, Mann-Whitney U test) higher than the 3 *year*^−1^ median amplitude (Figure 3B).

**Figure 3.**
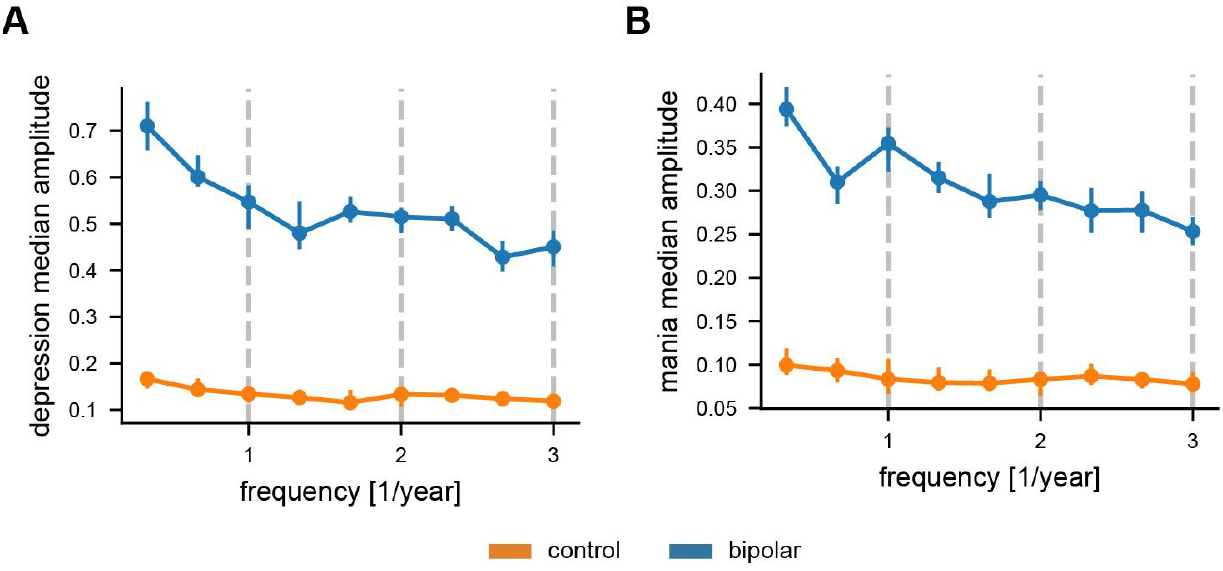
Frequency spectra of mood scales from a cohort of individuals with BD show year scale fluctuations. **(A)** Median Fourier amplitudes of PHQ-9 depression scale scores from 266 BD and 179 control participants measured every 2 months for at least two years. **(B)** Median Fourier amplitudes of ASRM mania scale scores from 273 BD and 178 control participants measured every 2 months for at least two years. Error bars are 68% confidence intervals from bootstrapping.

We conclude that hair cortisol and mood scales in BD show similar frequency distributions, with dominant low frequencies on the scale of a year that have higher amplitude than those on the scale of months.

### Mathematical model of the HPA axis in BD predicts enhanced year-scale fluctuations

In one of our prior studies (Maimon et al. 2020), a Fourier spectrum constructed from hair cortisol levels measured from control participants was similar to that in Figure 2B-C, with dominant low frequencies on the scale of a year. We showed that these frequencies cannot be explained by the classical mechanism of the HPA axis. The classical mechanism works on the timescale of the hormone lifetime, about an hour, and thus can not show timescales of months and years. We then showed that a recent mathematical model of the HPA axis (Karin et al. 2020; Tendler et al. 2021) that considers the temporal changes in the size of the HPA glands over months (Figure 4A) is able to reproduce the frequency spectrum of control participants. Stress inputs that vary from day to day, modeled as white noise, cause the glands to grow and shrink on the timescale of many months, providing the observed Fourier spectrum. The HPA system modulates the typical flat Fourier spectrum of the noisy input (Figure 4C) and amplifies low frequencies (Figure 4E) due to gland changes.

**Figure 4.**
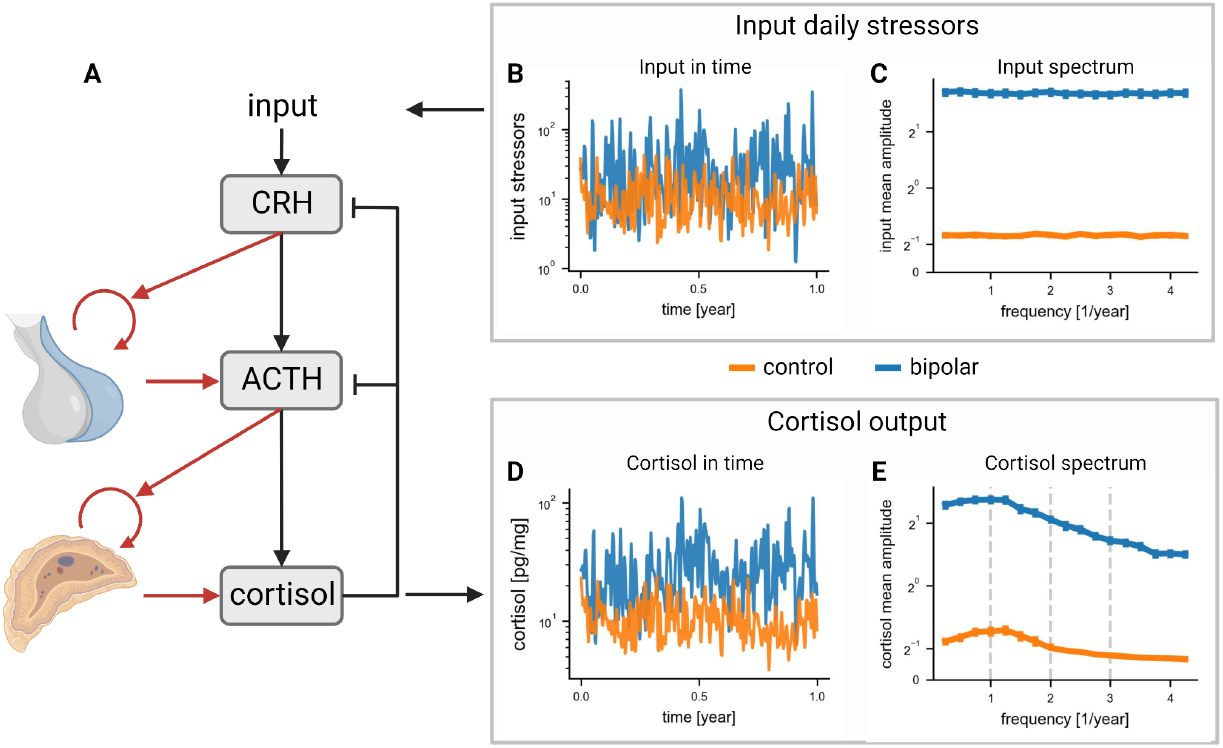
HPA model with gland mass changes predicts enhanced year-scale fluctuations in BD in agreement with the experimental measurements. **(A)** HPA axis circuit with gland mass dynamics. Red arrows represent growth-factor activities of hormones on the pituitary ACTH-secreting cells and the adrenal cortex. **(B)** One realization of simulated noisy daily stress inputs in time for BD (blue) and control (orange). **(C)** A flat Fourier spectrum of simulated noisy inputs shown in (B). **(D)** The simulated response of cortisol to the stress inputs shown in (B) over a year, where a baseline cortisol level was set to be the mean of the first segment hair data 10 pg/mg **(E)** Fourier spectrum of simulated cortisol shown in (D). The three gray dashed lines mark the frequencies that could be measured in the hair cortisol experiment.

Here we extend this mathematical model to identify a possible reason why the BD group exhibits a 4-fold increase in low frequency amplitudes compared to controls. We propose that the daily stress inputs to the hypothalamus in people with BD are higher than in the control population. We modeled this as a white noise input to the HPA equations which has a larger amplitude than in controls (Figure 4B-C).

We find, using simulations of the mathematical model, that the BD cortisol spectrum can be obtained by providing daily stress noise that is 4-fold larger than the noise needed to obtain the Fourier spectrum of control participants (Figure 4D-E). We thus conclude that a possible explanation of the enhanced slow cortisol fluctuations in BD participants may be larger day-to-day fluctuations in their stress inputs to the HPA axis.

### Hypothesis for a pathophysiological mechanism for the timescales of BD

Taking the present findings together, we propose a mechanism that could explain the timescales of the observed mood phenotypes in BD based on the HPA axis. This is built on decades of research showing the connection between the HPA axis and BD (A. H. Young and Juruena 2021; Belvederi Murri et al. 2016). The new aspect of the proposed mechanism is the ability of the HPA axis to generate fluctuations in cortisol on the time-scale of many months (Maimon et al. 2020) due to changes in the effective mass of the glands (Karin et al. 2020) (Figure 4A).

The basic premise of the proposed mechanism is that individuals susceptible to BD have two neurobiological traits (Figure 5A): a susceptibility in which high cortisol levels over weeks can trigger mood episodes, and emotional reactivity in which individuals generate larger daily stress inputs to the HPA axis in response to life events than the typical population. The HPA axis amplifies these enhanced inputs to generate large cortisol fluctuations over months that can trigger mania and depression.

**Figure 5.**
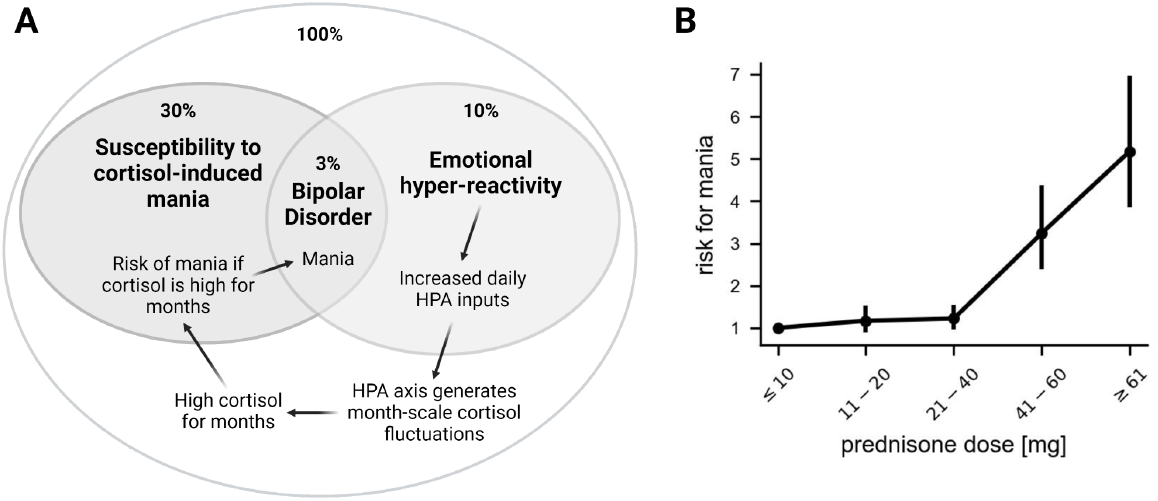
Hypothesis for a pathophysiological mechanism for the timescales of BD. **(A)** Overview of the present mechanism for bipolar disorder. Bipolar disorder can occur in individuals with both cortisol-induced mania (and depression) susceptibility and emotional hyper-reactivity. Emotional hyper-reactivity causes higher HPA inputs that fluctuate over hours and days. The HPA axis transforms these into months-scale fluctuations because of gland mass changes. High cortisol over months increases the probability of mania and depression in individuals with susceptibility to cortisol induced mood episodes. Percentages are estimates of the fraction of the population with each trait (see SI). **(B)** Risk of mania in a population (n=372,696) taking glucocorticoid steroids representing 89,298 years of glucocorticoid exposure compared to a control population. Normal cortisol baseline is equivalent to about 4 mg prednisone. Adapted from the data of (Laurence Fardet, Petersen, and Nazareth 2012), error bars are 95% CI.

These two neurobiological traits are supported by multiple lines of evidence. Susceptibility to cortisol-induced mania is found in studies on long-term use of glucocorticoid steroids, which are cortisol analogues. According to a large-scale study, people treated with glucocorticoids have a 2-fold higher risk of developing depression and a 4-5-fold higher risk of developing mania compared with people unexposed to glucocorticoids (Laurence Fardet, Petersen, and Nazareth 2012). The effect is dose-dependent (Figure 5B), with an apparent threshold of about 40 mg prednisone, equivalent to about 10 times the normal level of cortisol (Caetano and Malchoff 2022).

As described in the introduction, a similar effect is observed in Cushing’s syndrome in which cortisol levels are elevated due to a hormone-secreting tumor. Mania or hypomania is reported in about 30% of Cushing’s syndrome patients and depression in about 50-60% of the patients (Pivonello et al. 2015). In both Cushing’s syndrome and steroid treatment there is evidence that mania tends to occur earlier than depression (Pivonello et al. 2015; Judd et al. 2014).

Along with susceptibility to glucocorticoid-induced mood episodes, our proposed mechanism also requires special conditions which promote prolonged excess cortisol that can cross the mania and depression threshold. One such condition is a neuropsychological trait called emotional hyper-reactivity (Gross 1998), which is the generation of stronger-than-typical emotional responses to stimuli and is often reported in bipolar disorder (Dargél et al. 2020; Sperry and Kwapil 2022). We operationalize emotional hyper-reactivity by defining it as greater hypothalamic sensitivity leading to more pronounced input signals from daily stress than those of the typical population. A related variable is deregulation of circadian cycles as evident by dysregulated melatonin dynamics in BD. Circadian rhythm is an important input to the HPA axis. Both emotional reactivity and circadian dysregulation have fast-scale dynamics (hours-day) that can, in our model, excite slow timescale fluctuations (months) along the lines of Figure 4.

The heart of the proposed mechanism is the ability of the HPA axis to generate months-scale fluctuations of cortisol. This timescale of months is provided by the growth and shrinkage of the pituitary and adrenal functional masses. This months-scale property of the HPA axis converts input stresses that fluctuate rapidly over hours and days to large months-scale fluctuations. We posit that these months-scale fluctuations are larger in those with emotional hyper-reactivity because of their enhanced daily stress inputs. This coincides with the hair cortisol fluctuations measured here. As a result of prolonged periods of high cortisol, there is an increased probability of triggering mood episodes.

The prevalence of BD in the general population is about 2%. This may correspond to the intersection of the two traits mentioned above. Cortisol-induced mania appears in about 30% of Cushing’s patients, so that susceptibility to cortisol-induced mania may characterize about 30% of the general population. Emotional hyper-reactivity appears in at least 10% of the general population (see SI). If these traits are independent, one may expect about 3% to have both (Figure 5A).

## Discussion

We present longitudinal hair cortisol measurements from participants with BD combined with mathematical modeling to propose a mechanism for mood fluctuations in BD. Participants with BD showed higher mean hair cortisol and enhanced cortisol fluctuations on the year-timescale as compared to a control population. Hair cortisol in the proximal 2cm segment correlated with mood scales within the BD cohort. The frequency spectra of cortisol is similar to that of mood scales from a previous longitudinal BD cohort. The enhanced year-scale fluctuations can be explained by using a model of the HPA axis based on fluctuations in gland masses on the timescale of months. We discuss a mechanism for the timescales of BD based on these results.

This study aligns with the view that longitudinal measurements are important for studying BD, an inherently complex and dynamic disease (McInnis et al. 2022). Studies that explore a single time point can assess the differences in mean between populations, such as higher mean cortisol in BD (Belvederi Murri et al. 2016; Coello et al. 2019; Streit et al. 2016), also found here. Single time point studies, however, cannot address questions like the amplitude of 12-month or 6-month cortisol fluctuations. The present data indicate that such months-scale cortisol fluctuations are larger in BD than in control populations.

To measure cortisol we used hair, because it offers advantages over other assays, namely blood, saliva and urine measurements (D’Anna-Hernandez et al. 2011; Davenport et al. 2006; Mayer et al. 2019; Sauvé et al. 2007). Cortisol accumulates passively in the hair, and protocols exist to measure hair cortisol. A 2cm hair sample represents about 2 months of growth, offering a window into the mean cortisol over the last two months (D’Anna-Hernandez et al. 2011; Smy et al. 2016). For example, hair cortisol reports accurately on cortisol dynamics over months in patients with Cushing’s syndrome (Thomson et al. 2010). Averaging cortisol over 2 months bypasses the sensitivity of blood or saliva cortisol tests to circadian rhythm, the menstrual cycle and acute stresses at the time of the test. Hair is also easy to collect and store.

We used 12cm of hair, and each participant filled out mood scales when hair was collected. We could thus correlate mood with the proximal 2cm of hair that represents 2 months of growth. Hair cortisol correlates well with depression scales in the BD cohort. Since no participant had high mania scale scores, we could not deduce association between hair cortisol and mania scales. Further study can collect shorter hair segments repeatedly along with mood scales, in order to test the temporal correlation of cortisol and mood in each individual over time.

The enhanced months-scale fluctuations in cortisol offer a possible origin, at least in some BD patients, for the timescales of mood swings. Enhanced month-scale fluctuations are relevant to BD when the following is true. First, the individual needs to produce larger daily stress inputs than is typical in the population, in terms of hypothalamic CRH secretion to the HPA axis, due to their CNS makeup and their life events. Such enhanced input causes the slowly-varying HPA glands to fluctuate in mass, and to produce enhanced month-scale cortisol fluctuations. Second, the individual needs to be susceptible to cortisol induced mood episodes. Such susceptibility may be analogous to that seen in individuals treated with glucocorticoids that respond with BD-like symptoms. For the BD patients that meet these two conditions, a strategy that reduces cortisol and its fluctuations might be therapeutic.

Other physiological mechanisms that can in principle underlie the months-timescale of BD might include epigenetic modifications, such as DNA methylation and histone acetylation, which can have timescales of months, and may conceivably affect mood episodes. In contrast, other biological processes such as neurotransmitter function and gene expression are unlikely to accommodate the observed time scale because they typically work on the timescale of hours to days.

The mathematical model for the HPA axis used here has been tested in several contexts, such as hormone seasonality (Tendler et al. 2021), HPA function in addiction (Karin, Raz, and Alon 2021), HPA recovery from prolonged stress (Karin et al. 2020), and cortisol variation in healthy controls (Maimon et al. 2020). Therefore, it seems to be a reasonable description of the slow timescale of the HPA axis.

### Limitations of the Study

This study involved a sample of only 59 control participants and 26 BD participants from one country, primarily female. Future work can enlarge sample size and sample additional populations, which is important given the large person-to-person variability in cortisol. Use of other methods to measure cortisol, such as mass spectrometry of hair samples, or blood urine or saliva cortisol assays, can test the validity of the results. Use of multiple hair samples from the same individual over time can extend the study period beyond 12 months.

In summary, we present data on hair cortisol in BD participants and find dominant low frequencies of a year. We find a similar frequency spectrum in mood scales from a large cohort of people with BD. The frequency spectrum is explained by a mathematical model of the HPA axis due to the slow growth and shrinkage of the gland functional mass. Taken together, this data suggests a three way correspondence of cortisol, mood scales and HPA physiology. It therefore suggests a mechanism for the timescales of BD based on the HPA axis amplification of enhanced daily stress inputs in people where cortisol can induce mood episodes. We hope that these findings will lead to further exploration of the timescales of BD and their physiological origin, in order to better understand the etiology of this disorder.

## Materials and Methods

### Ethics statement

The study protocol was approved by the Helsinki committee of Be’er Ya’akov mental hospital (study code: 613) and by the Review Board of the Weizmann Institute of Science (study code: 1312-1). Written informed consent was obtained from all participants.

### Participants

#### Control group

Healthy participants (N=71, females=55, average age=28 ± 5 years) were recruited using social media. Inclusion criteria were age 18-65 years, with at least 12 centimeters of natural hair with no cosmetic treatment such as dying or perming (Cooper 2015). Exclusion criteria were diagnosis of a mental, psychological or endocrine disorder, consumption of steroids, psychiatric drugs or other drugs that might affect the endocrine system during the year prior to participation, and pregnancy in the year before participating. Oral contraceptives were allowed as long as there was no change in prescription during the year prior to participation. Each subject was asked to complete a personal information questionnaire prior to the collection of a hair sample. The study was anonymous; each hair sample received a serial number.

The 71 control participants include 59 participants from a previous study (Maimon et al. 2020). Of these, 21 participants returned to provide a new 12 cm hair sample for the present study, and this data was used. For the remaining 38 participants from the previous study, the previous data was used.

#### Bipolar disorder group

Participants diagnosed with BD (N=28) were recruited from Be’er Ya’akov mental hospital and via ads approved by the Weizmann Institute IRB (study code 1312-1). Inclusion criteria were diagnosis of BD (DSM III-V), age 18-65, stable medical treatment in the last 2 months or without medical treatment, with at least 12 centimeters of natural hair with no cosmetic treatment such as dying or perming (Cooper 2015). Exclusion criteria: substance use disorder, diagnosis of PTSD (due to reports of lower resting levels of circulating cortisol, and from consistent evidence of heightened HPA axis response to challenge tests in chronic PTSD (de Kloet et al. 2006; Yehuda 2002) or schizophrenia, steroid treatment in the 6 months prior to participation, surgery or brain stimulation in the 6 months prior to participation and pregnancy in the year before participating. Oral contraceptives were allowed as long as there was no change in prescription during the year prior to participation. The study was anonymous; each hair sample received a serial number, case report form (CRF) was coded with initials and a serial number.

### Hair Cortisol Measurements

We used the method of (Maimon et al. 2020). A lock of hair (a pencil-width group of about 100 hair strands) from the vertex posterior area (Cooper 2015; Sauvé et al. 2007) of the head was tied with a thread and cut with fine scissors as close to the scalp as possible. We estimate that hair was cut at 0.8±0.1 cm from the scalp, in agreement with (Cooper 2015). Cut samples were kept in aluminum foil; the tied thread marked the proximal end. Samples were kept in the laboratory at room temperature before analysis. We adapted a protocol by (Schonblum et al. 2018) for the extraction and measurement of hair cortisol. The first 12 centimeters of each hair sample, starting from the proximal end, were segmented to six 2 cm segments (segments of 1 cm dropped below the detection threshold too often and thus 2 cm segments were used). The segments were placed in vials (Fisherbrand, 21×70 mm) and washed twice with 5 ml isopropanol while mixing on an orbital rotator for 3 minutes. Isopropanol was then decanted and the open vials were left in a chemical hood to dry overnight. Then, 2 ml of methanol was added to each vial, sonicated for 60 min, and incubated overnight (approximately 20 hours) at 50°C while shaking. The following day, all the methanol was transferred to 2 ml Eppendorf tubes and centrifuged for 10 minutes at 4°C. Methanol (1.5 ml) from each tube was transferred to a glass vial (Falcon, 12×75 mm) and evaporated under a stream of nitrogen at 45 °C. Samples were reconstituted in 10% methanol and 90% assay buffer provided by the kit manufacturer and cortisol was quantified using competitive Enzyme-Linked Immunosorbent Assays (ELISA; Salimetrics Europe, Newmarket, cat.no1-3002-5 for cortisol, UK). Reported antibody cross-reactivity was 19.2% with dexamethasone, and less than 1% with 15 other tested steroids. Linearity was observed between 30-70 mg of hair, hence we used 30-70 mg of hair to measure cortisol. The assay detection threshold was 112 pg as specified by the manufacturer. All 6 segments from the 12 cm sample of hair from each participant were analyzed in the same batch of washing, sonication, extraction and ELISA plate. To control for inter-assay variation, we generated a standard curve for each plate, consisting of 6 known concentrations of cortisol supplied by the kit manufacturer assayed in 6 wells. To estimate the inter-batch variation, we assayed multiple standard hair samples. Each standard sample was taken from a large, well-mixed, sample of hair collected from a single individual. The coefficient of variation (CV=SD/mean) of 13 standard samples measured on 2 different days was 14%.

We included in this study the 91 participants (63 controls, 28 with bipolar disorder) that had all 6 cortisol measurements above detection threshold.

### Hair cortisol time series analysis

We used the method of (Maimon et al. 2020) to correct for the decline along the hair and obtain normalized cortisol levels from the 6 hair segments. We calculated the mean, standard deviation (SD), coefficient of variation (CV=SD/mean) and the Fourier spectrum for each participant as described in (Maimon et al. 2020). We removed participants with average cortisol or cortisol CV that was higher by more than three SDs than the population mean. Specifically, we removed one control participant and two BD participants with cortisol exceeding 240 pg/ml, and 3 control participants with CV that exceeded 1.1. The analysis thus included 59 healthy controls and 26 BD patients. 13 BD participants had repeat measurements spaced by 2 months or more. When comparing raw cortisol, we averaged the first and second repeats per segment. When assessing cortisol temporal fluctuations we averaged the Fourier amplitudes of the first and second repeats for each of these 13 participants.

### Psychological Measurements

Interviews were conducted by a graduate student with training and supervision of a psychiatrist. Each participant was interviewed to collect medical history, medication treatment and mood evaluation. The interview included a clinical interview and supervised taking of the Hamilton Depression Rating Scale (HDRS) (Hamilton 1960), together with a self-report using the Beck Depression Inventory (BDI) (Beck et al. 1961). Mania was rated using the Young Mania Rating Scale (YMRS) (R. C. Young et al. 1978). Anxiety was measured using the Hamilton Anxiety Rating Scale (HAM-A) (Hamilton 1959), general functioning was assessed using the Clinical Global Impressions inventory (CGI) (Guy 1976) and the Global Assessment of Functioning scale (GAF) (Aas 2010). Interviews were conducted with training and supervision of a psychiatrist.

### Association of hair cortisol and mood scales

One participant out of the 26 BD participants didn’t fill mood questionnaires. Thus, a total of 25 participants were analyzed. 13 participants returned for a second measurement. They were thus tested twice at intervals longer than 2 months, to provide a total of 38 hair samples with 38 corresponding sets of psychological scores. Sample size meets the requirements for detecting correlation of 0.45 with power = 0.8 and alpha = 0.05. We used log cortisol to adjust for the variation in cortisol between individuals (Maimon et al. 2020). We calculated partial Pearson correlations in order to adjust for participant’s age, gender, family status, education and medication. The null hypothesis was that cortisol does not correlate positively with scale scores. Hence we used a one tailed statistical test to evaluate significance. Statistical analysis was conducted using Python Scipy and pingouin packages (Virtanen et al. 2020; Vallat 2018).

### Participants from Prechter Cohort

Participants were enrolled in the Prechter Bipolar Longitudinal Cohort (McInnis et al. 2018), an observational cohort study gathering phenotypic and biological data, at the University of Michigan. We recruited participants into the cohort through advertisements on the web, in the newspaper, in an outpatient specialty psychiatric clinic, community mental health centers, community outreach events and in an inpatient psychiatric unit from 2006-2018. At study entry, BD and healthy controls were evaluated using the Diagnostic Interview for Genetic Studies (Nurnberger et al., 1994) and a best estimate process by at least two of the authors was used to confirm diagnosis using DSM-IV criteria. Participants were excluded if they had active/current substance abuse (per DSM-IV criteria) at enrollment or neurological disease. Healthy controls were included if they had no history of DSM-IV axis I psychiatric illness and no family history of psychiatric diagnosis. This study was approved by the University of Michigan IRB and written informed consent was obtained and incentive payment for their participation in the study was provided.

We analyzed data collected from bipolar I and control individuals (Table 2) followed prospectively for at least 4 years in the Prechter Longitudinal Study of Bipolar Disorder at the University of Michigan (McInnis et al. 2018; Langenecker et al. 2010). The Altman Self-Reported Mania (ASRM) (Altman et al. 1997) and the Patient Health Questionnaire for Depression (PHQ9) (Kroenke, Spitzer, and Williams 2001) scales were completed at 2 month intervals. The UM IRB approved recruitment, assessment, and research procedures (HUM606).

**Table 1.** Demographic and clinical characteristics of participants with bipolar disorder.

|  |  |
| --- | --- |
| **Total BD group** | N=26 |
| Male; n (%) | 5 (19) |
| Age (year); median (IQR) | 32 (13) |
| **family status** |  |
| single; n (%) | 18 (69) |
| divorced; n (%) | 5 (19) |
| married; n (%) | 3 (12) |
| **Number of children** |  |
| 0; n (%) | 14 (54) |
| 1; n (%) | 5 (19) |
| 2; n (%) | 4 (15) |
| missing | 3 (12) |
| **years of education** |  |
| less than 9; n (%) | 0 (0) |
| 9 to 12; n (%) | 5 (19) |
| more than 12; n (%) | 21 (81) |
| **Medicated** |  |
| medicated; n (%) | 14 (54) |
| irregular; n (%) | 2 (8) |
| unmedicated; n (%) | 10 (38) |
| **Medications** | N=16 |
| mood stabilizer; n (%) | 15 (94) |
| antipsychotics; n (%) | 10 (62.5) |
| antidepressants; n (%) | 10 (62.5) |

**Table 2.** Demographic characteristics of participants from the Prechter cohort. * Participants with at least two years of consecutive mood measurements

|  | Participants with depression scores* |  | Participants with mania scores* |  |
| --- | --- | --- | --- | --- |
|  | Bipolar (N=266) | Control (N=179) | Bipolar (N=273) | Control (N=178) |
| **Gender** |  |  |  |  |
| Male; n (%) | 78 (29.3) | 61 (34) | 79 (28.9) | 61 (34.3) |
| Female; n (%) | 187 (70.3) | 118 (66) | 193 (70.7) | 117 (65.7) |
| Other; n (%) | 1 (0.4) | 0 (0) | 1 (0.4) | 0 (0) |
| **Age** (year); median (IQR) | 51.5 (22) | 43 (26) | 52 (22) | 43 (26) |
| **Race** |  |  |  |  |
| White; n (%) | 242 (91) | 138 (77.1) | 248 (90.8) | 139 (78.1) |
| Black or African-American; n (%) | 5 (1.9) | 15 (8.4) | 7 (2.6) | 15 (8.4) |
| Asian; n (%) | 3 (1.1) | 16 (8.9) | 3 (1.1) | 16 (9) |
| American Indian/Alaska Native; n (%) | 0 (0) | 1 (0.5) | 0 (0) | 1 (0.6) |
| More Than One Race; n (%) | 14 (5.3) | 7 (4) | 13 (4.8) | 5 (2.8) |
| Unknown or not reported; n (%) | 2 (0.7) | 2 (1.1) | 2 (0.7) | 2 (1.1) |
| **Ethnicity** |  |  |  |  |
| Hispanic or Latino; n (%) | 10 (3.7) | 11 (6.1) | 11 (4) | 10 (5.6) |
| Not Hispanic or Latino; n (%) | 255 (95.9) | 165 (92.2) | 261 (95.6) | 165 (92.7) |
| Unknown or not reported; n (%) | 1 (0.4) | 3 (1.7) | 1 (0.4) | 3 (1.7) |

### HPA model

We used a mathematical model for the HPA axis that incorporates the pituitary and adrenal gland mass dynamics (Karin et al. 2020; Tendler et al. 2021). To simulate noisy daily stress input, we generated a signal that is a sum of 360 sine functions, each with different frequency (ranging from 1 per 4 years up to 1 per 4 days) and a random phase (Figure 4B-C). To analyze cortisol response to noisy stressors, we generated 1,000 simulations with such input signals. We used a burn-in period of one year and then computed the average Fourier amplitudes of cortisol over 4 additional years.

## Acknowledgments

This research was supported by Heinz C Prechter Bipolar Research Program at the University of Michigan Eisenberg Family Depression Center and the Richard Tam Foundation. With gratitude, we acknowledge the Bipolar Longitudinal Research participants and thank the research team lead for collection, stewardship and sharing of the data used in this publication.

We would like to thank Dr. Eric Simon of the Research Innovation Core at the University of Michigan’s Eisenberg Family Depression Center for reviewing and editing this manuscript.

Figures 4 and 5 were created with BioRender.com

## Resource Availability

Further information and requests for resources should be directed to and will be fulfilled by the Lead Contact, Uri Alon.

## Materials Availability

This study did not generate new unique reagents.

