## Supplementary Information for "Longitudinal hair cortisol in bipolar disorder and a mechanism based on HPA dynamics"

**Estimation of hyper-reactivity prevalence**

We didn’t find a previous estimate for the prevalence of hyper-reactivity in the population. To estimate this prevalence we summarized the prevalences of all the pathological conditions that are well-associated with hyper-reactivity, which total to 38% (Supporting information, Table S1). We then accounted for comorbidities (Supporting information, Table S2) to find an estimated hyper-reactivity prevalence of 11%. This is an underestimate in the sense that we don't take into account individuals without pathology.

| **Disorder** | **Type of Hyperreactivity** | **Prevalence in Population** |
| --- | --- | --- |
| Bipolar^1,2^ | Emotional Reactivity General  problematic emotional-regulation goals | 1-2%^3^ |
| Borderline Personality^1, 2^ | the duration of emotion is both too long and too short | 1-3%^4,5^ |
| Social Anxiety Disorder^2^ | hyperreactivity to specific negative emotions | 12.1%^6^ |
| Major Depressive Disorder^2^ | both hyperreactivity and hyporeactivity | 4.7%^7^ |
| Specific Phobia^2^ | duration of negative emotion is too long | 7.2%^8^ |
| Intermittent Explosive Disorder^2^ | emotions occur too frequently | 0.8%^9^ |
| Autism Spectrum Disorder^2^ | emotions may occur both too frequently and too infrequently | 0.76%^10^ |
| ADHD^2^ | problematic emotional regulation strategies | 9.34%^11^ |

**Table S1. Prevalence of hyper-reactivity-associated diseases**

|  | Bipolar | Borderline Personality | Social Anxiety Disorder | Major Depressive Disorder | Specific Phobia | Intermittent Explosive Disorder | Autism Spectrum Disorder | ADHD |
| --- | --- | --- | --- | --- | --- | --- | --- | --- |
| Bipolar |  | 10-20%^12^ | 13.3%^13^ |  | 66.6%^14^ | 6.5%^15^ | 1.4%^16^ | 10-21%^17^ |
| Borderline Personality | 20%^12^ |  | 42.4^18^ | 49.1^19^ - 61%^18^ | 20.3^18^ |  | 3%^20^ | 16.1-38.1%^21^ |
| Social Anxiety Disorder | 3.5-21%^22^ |  |  | 35-70%^22^ | 14.1-60.8^22^ |  |  | 60-70%^22^ |
| Major Depressive Disorder |  | 1.8-25^23^ | 20-30%^22^ |  | 13.5%^24^ | 17%^25^ |  | 34%^26^ |
| Specific Phobia |  |  | 6.7%^27^ | 40.7%^28^ |  |  |  |  |
| Intermittent Explosive Disorder | 6.9-8.4^29^, 17.6^30^ | 48.2%^31^ | 23.4^30^ | 26.4%^30^,  19-22.3%^29^, 21.5^31^ | 26.9^30^ |  |  | 12.7^32^ |
| Autism Spectrum Disorder | 6-40%^33^ | 4%^20^ | 20%^34^ | 37%^34^ | 14%^35^ | 16.67%^36^ |  | 38%^37^ |
| ADHD | 5-20%^17^ | 18-27%^38^ | 10-45% (*GAD)^39^ | 25%^40^ | 8.8^40^ | 24.5^32^ | 30%^37^ |  |

**Table S2. Comorbidities between hyper-reactivity-associated diseases**

**Resources**
